## Supplemental Material for "Parabolic avalanche scaling in the synchronization of cortical cell assemblies"

*bioRxiv*

### Supplementary Information

*Elliott Capek[1*], Tiago L. Ribeiro[1*], Patrick Kells[1], Keshav Srinivasan[1], Stephanie R. Miller[1], Elias Geist[1], Mitchell Victor[1], Ali Vakili[1], Sinisa Pajevic[1], Dante R. Chialvo[2], & Dietmar Plenz[1†]*

[1]Section on Critical Brain Dynamics, National Institute of Mental Health, Bethesda, MD, USA

[2]CEMSC3, Escuela de Ciencia y Tecnologia, UNSAM, San Martín, P. Buenos Aires, Argentina

*Co-first authorship

† *Correspondence*

Figures: 18 Supplementary Figures

Running Title: Parabolic avalanches in cortical synchronization

Correspondence: Dietmar Plenz, Ph.D., Section on Critical Brain Dynamics, National Institute of Mental Health, Porter Neuroscience Research Center, Rm 3A-1000, 35 Convent Drive, Bethesda, MD 20892. office: (301) 402-2249, fax: (301) 480-7480,

**Supplementary Figure S1**

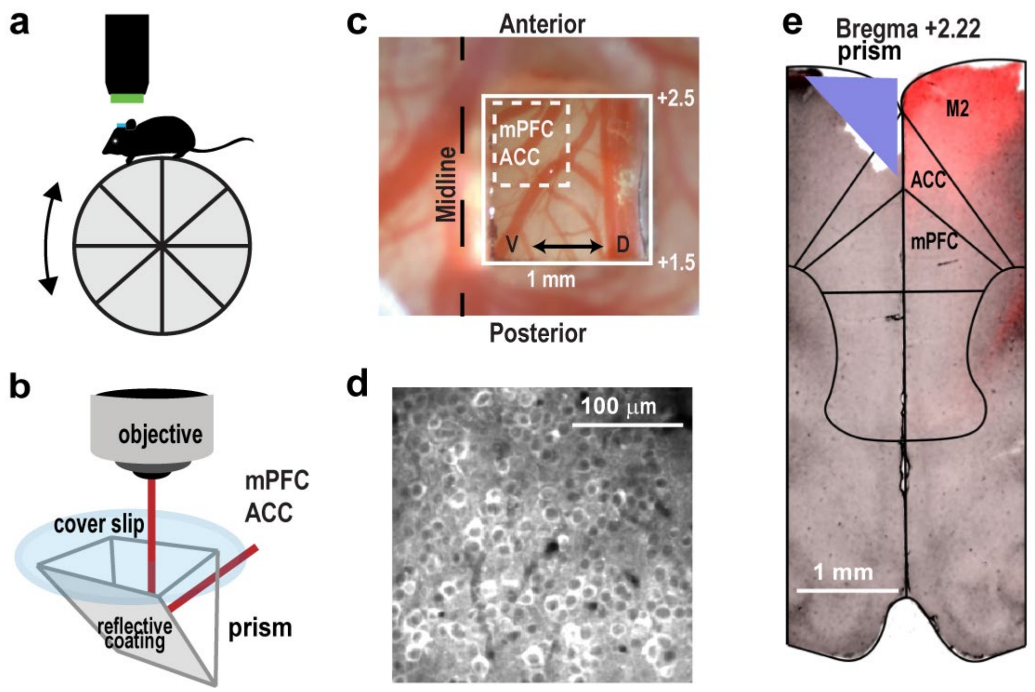


**Supplementary Figure S1| Overview of prism recording conditions and anatomy.** (**a**) Sketch of recording ongoing activity using 2PI in head-fixed mouse on a freely moving wheel. (**b**) A chronically implanted prism allows for imaging neuronal activity across the midline in the contralateral mPFC and ACC. (**c**) Dorsal view of cortex with implanted microprism (*white square*; bregma +1.5 - +2.5) in right hemisphere allowing for imaging the contralateral, medial cortex across the fissure (*midline*; light microscopic image). Within the prism, the broken square indicates the 450 µm x 450 µm recording region targeting the contralateral mPFC/ACC. *V*: ventral. *D*: dorsal. (**d**) jrGECO1a expressing neurons in contralateral mPFC/ACC (~150 µm from surface; average 2PI image for ROI detection). (**e**) Anatomical recovery of prism position in ipsilateral frontal cortex (*triangle*) and corresponding large-field jrGECO1a fluorescence (*red*) in contralateral cortical regions (coronal brain section at bregma +2.22).

#### Supplementary Figure S2

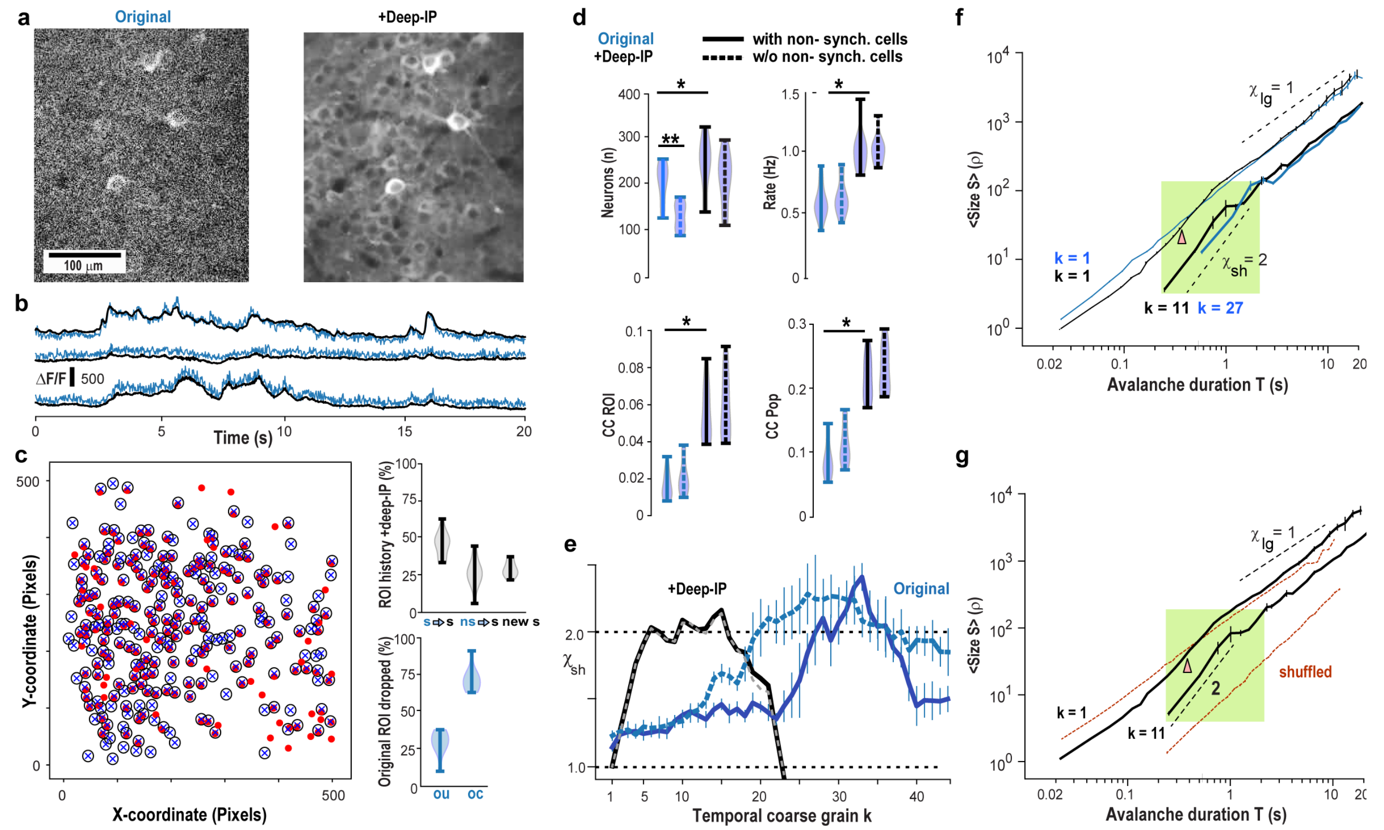


**Supplementary Figure S2 | Improvement in prism-based 2-PI of ongoing neuronal activity by adding deep-IP and its effect on avalanche scaling. (a**) Example of a single 2PI frame (22 ms) before (*original*) and after including spatiotemporal deep-interpolation (+*Deep-IP*; mPFC; prism; red-shifted jRGECO1a, single mouse). (**b**) Deep-IP reduces noise-fluctuations in relative fluorescent change ΔF/F (3 example ROIs, shifted for clarity). (**c**) Example of ROI identification in the original Field-of-View (*square*) and after applying Deep-IP. *Filled red circle*: original, population correlated ROI. *X*: +Deep-IP. *Open circle/X*: Original ROI included after deep-IP (jrGECO1a, single mouse) *Top right*: The population of correlated neurons in +Deep-IP originate from originally synchronized (s→ s, *left*) and non-synchronized neurons (ns → s, *middle*), as well as newly identified synchronized neurons (new s, *right*). *Bottom right*: The small fraction of neurons in the original analysis not recognized anymore after +Deep-IP (set to 100%) are proportionally drawn from originally uncorrelated (*ou*) and correlated (*oc*) neurons. (**d**) Summary statistics in change in number of neurons (*Neurons*), mean spike rate (*Rate*), average cross correlation between neurons (*CC ROI*), and between single neurons and the remaining population of neurons (*CC Pop*). Data describe neuronal statistics for the original analysis (*Original*), with Deep-IP added (+*Deep-IP*) and when non-synch cells are included (*solid lines*) or removed (*broken lines*). Note that after adding Deep-IP, the prevalence and impact of non-synchronized neurons is greatly diminished, while rate and CC values are significantly increased. A minimal z-score ($p < 0.001$) in the correlation with population activity was used to distinquish synchronized from non-synchronized neurons. (**e**) Deep-IP greatly improves the temporal resolution (low k) at which $\chi_{sh}$ is obtained. Summary of $\chi_{sh}$ as a function of $k$ (For legend see *d*). (**f**) $\chi_{sh} = 2$ holds for avalanche durations ~0.2 – ~2 s at $k = 11$ (*green shaded region*) and $k = 27$. Note that after +DeepIP, $\chi_{sh} = \sim 2$ is approximated for avalanches of similar absolute duration now visible at $k = 1$ (*arrowhead*), which was not present in the original analysis. (**g**) Shuffle comparison for k = 1 and k = 11. As shown in *f*, $\chi_{sh} = 2$ holds for avalanche durations T = ~0.2 – 1.3 s at $k = 11$ (*green region*), which is abolished by temporal shuffling (*red*). $k = 1$ shown for comparison. (*c – g*, averages over n = 5 mice; jRGECO1a). Values are mean ± SD.

#### Supplementary Figure S3

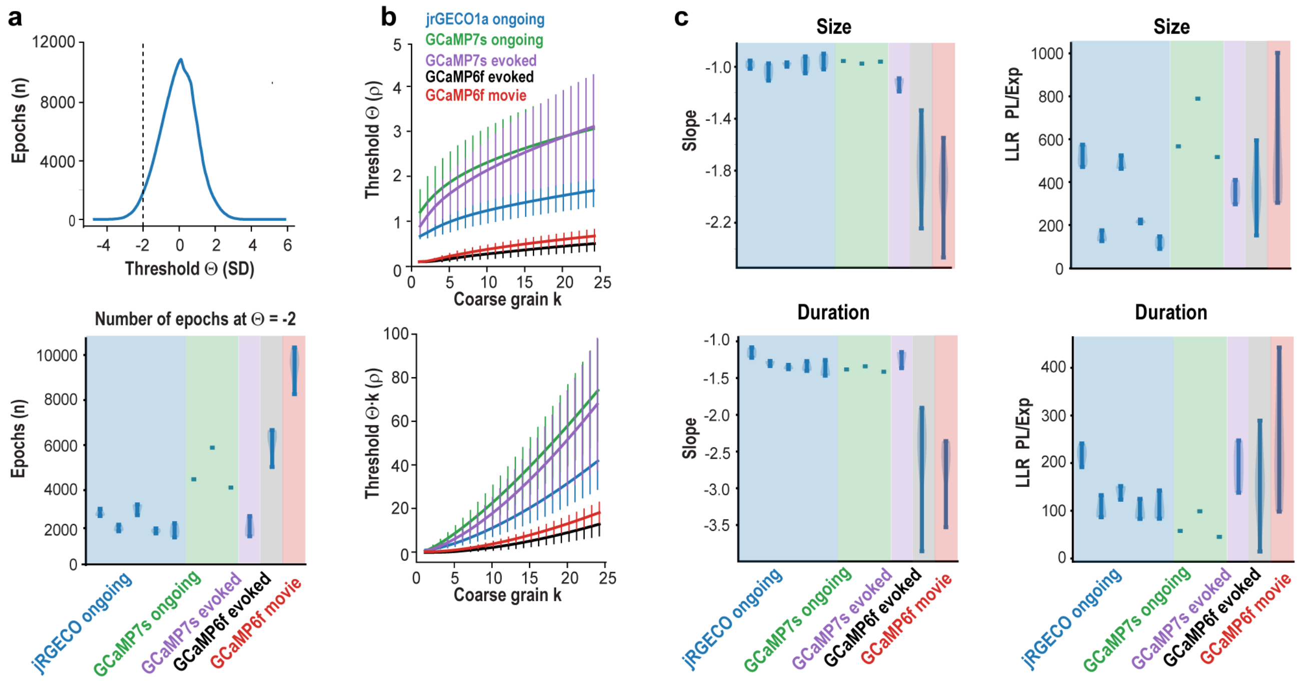


**Supplementary Figure S3 | Summary statistics on number of epochs, changes to threshold Θ, and LLR. (a**) *Top*: Number of epochs distribute approximately lognormal and were plotted as a function of z-score (jRGECO1a; example of single mouse). All results presented in the main text were obtained at Θ = -2 SD (*broken line*). *Bottom*: Summary of number of epochs obtained at Θ = -2 SD for all experimental groups. Individual mice were averaged per group (jRGECO ongoing, +Deep-IP; GCaMP7s ongoing, non-synch ROI removed; GCaMP7s drifting gratings, +Deep-IP). Averages over all mice are shown for the Allen Institute data set (GCaMP6f drifting, movie). (**b**) The requirement of Θ(-2SD) after temporal coarse graining leads to a monotonic increase in absolute Θ (*top*; mean ± SD; per experimental condition). This increases the synchrony requirement $\Sigma\rho/\Delta t \geq \Theta(k)$ with $k$, while relaxing the continuous epoch criterion. For comparison, the total amount of subthreshold activity removed from the analysis also increases monotonically (*bottom*). Average over all mice and experiments per experimental condition. Mean ± SD. (**c**) *Left*: Power law slopes for size and duration distributions at original frame rate ($k = 1$). Note steeper slopes for GCaMP6f, which exhibits less sensitivity to action potential firing compared to GCaMP7s and jRGECO. *Right*: Corresponding summary statistics for the LLR when comparing a power law (*PL*) against an exponential (*Exp*) distribution. Consistently positive LLR ratios demonstrate that distributions in epoch size and duration demonstrate epochs fulfill the definition of neuronal avalanches. *a*, *c* violin plots.

#### Supplementary Figure S4

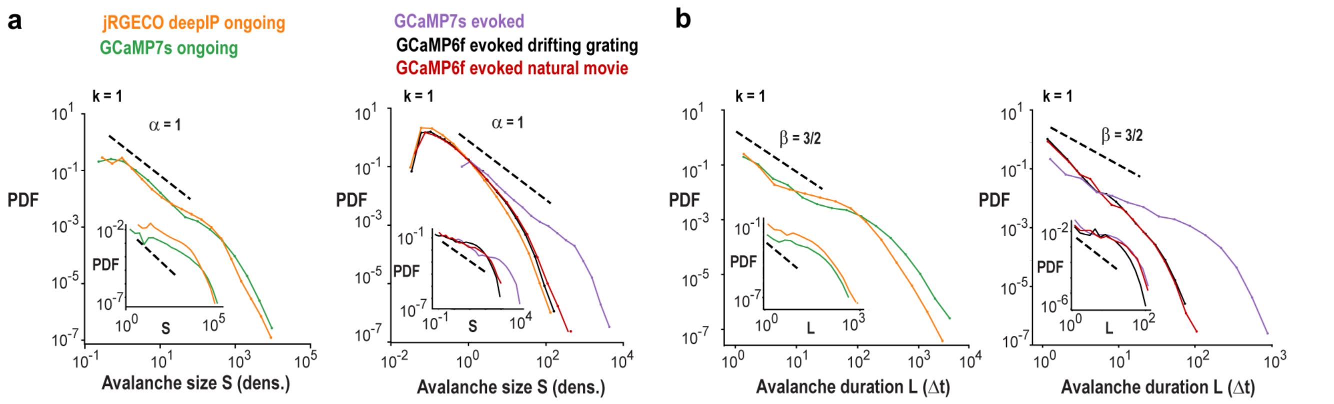


**Supplementary Figure S4 | Average size and duration distributions at frame rate resolution ($k$ = 1) and $\chi_{sh}(k) \cong 2$ for all experimental conditions.** (**a**) Size distribution for ongoing (*left*) and evoked (*right*) avalanches at $k$ = 1. *Insets*: Corresponding size distributions for $k$ at which $\chi_{sh}(k) = 2$. (**b**) Corresponding duration distributions. Averages over all mice for each condition. *Broken lines*: Guides to the eye.

**Supplementary Figure S5**

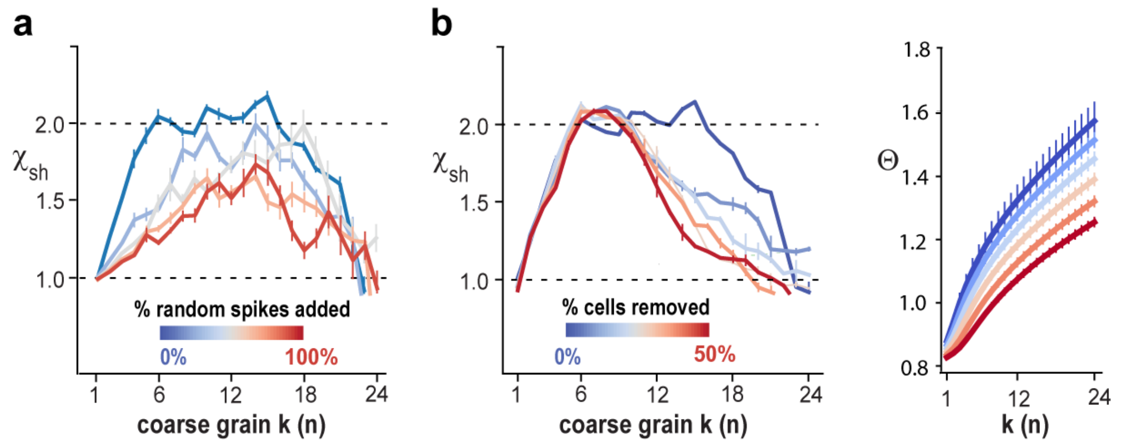


**Supplementary Figure S5 | Recovery of $\chi_{sh}$ = 2 depends on signal quality and is robust to small changes in subsampling fraction.** (**a**) The addition of random spikes shifts the maximal obtainable value of $\chi_{sh}$ towards higher temporal coarse graining (jrGECO1a; n = 5 repeats). Note that high randomization values eventually reduce the ability to recover $\chi_{sh}$ = 2. (**b**) Recovery of $\chi_{sh}$ = 2 is robust to up to 50% reduction in subsampling fraction *f*. *Left*: $\chi_{sh}$ (*k*) for different percentage of cells removed from the analysis. A reduction in *f* by half (50%) still allows for recovery of $\chi_{sh}$ = 2. *Right*: Change in $\Theta$ as a function of *k* and percentage of removed cells. $\Theta$ at which -2SD of the maximal number of avalanches is reached increases with *k* and decreases with *f*. The compensation originates from a corresponding overall decrease in $\Theta$ in identifying avalanches, i.e., higher sensitivity in utilizing instantaneous population activity (*right*). Deep-IP jRGECO ongoing activity used in all subpanels (n = 5 mice; cp. Fig. 1). *Black broken lines*: Reference for χ. Error bars: Bootstrapping with n = 10.

#### Supplementary Information S6

##### *Error estimates when deriving the parabolic scaling exponent using thresholding*

Starting with a generic inverted parabola, $y = Ax(D - x)$, with amplitude *A* and roots *0, D*, we next make the requirement that the avalanche size, *S*, i.e., area underneath the parabola, needs to scale quadratically with the avalanche duration, *D,* (Suppl. Fig. S6a)

$$S = \int_0^D Ax(D - x)\, dx \propto D^2. \quad (1)$$

Given $S = cD^2$, with *c* being a constant, and solving the integral in (1), we obtain $A = 6c/D$. Therefore, $y = 6cx(D - x)/D$ represents self-similar parabolic avalanches with duration *D* for any *c* in which size *S* scales quadratically with *D*.

We then proceed to apply a threshold Θ to this original parabolic shape, which changes both our perceived duration and size (Suppl. Fig. S6a). The thresholded duration, $D_\Theta$, will be given by the difference between the roots of the thresholded parabola

$$y_\Theta = y - \Theta = 6cx(D - x)/D - \Theta. \quad (2)$$

Solving for the roots, we have $6cx(D - x)/D - \Theta = 0 \rightarrow -6cx^2/D + 6cx - \Theta = 0$, with roots

$$x_{0+,-} = (D/2)\left(1 \pm \sqrt{1 - 2\Theta/3Dc}\right), \quad (3)$$

from which we obtain: $D_\Theta = x_{0+} - x_{0-}$,

$$D_\Theta = D\sqrt{1 - 2\Theta/3Dc}. \quad (4)$$

When using a *soft threshold*, the avalanche size is defined as the area of the parabola above the threshold, $S_\Theta = \int_{x_{0-}}^{x_{0+}} (6cx(D - x)/D - \Theta)\, dx$, where $x_{0+}$ and $x_{0-}$ are the roots of the original, non-thresholded parabola in (2). By solving the integral, we obtain

$$S_\Theta = -(2c/D)({x_{0+}}^3 - {x_{0-}}^3) + 3c({x_{0+}}^2 - {x_{0-}}^2) - t(x_{0+} - x_{0-}). \quad (5)$$

Using (3) and (4), we have

$$S_\Theta = -(cD^2/4)((1 + D_\Theta/D)^3 - (1 - D_\Theta/D)^3) + (3cD^2/4)((1 + D_\Theta/D)^2 - (1 - D_\Theta/D)^2) - \Theta D_\Theta$$

$$= (cD^2/4)\{(1 + D_\Theta/D)^2[3 - (1 + D_\Theta/D)] - (1 - D_\Theta/D)^2[3 - (1 - D_\Theta/D)]\} - \Theta D_\Theta$$

$$= (cD^2/4)\{[1 + 2D_\Theta/D + (D_\Theta/D)^2](2 - D_\Theta/D) - [1 - 2D_\Theta/D + (D_\Theta/D)^2](2 + D_\Theta/D)\} - \Theta D_\Theta$$

$$= (cD^2/4)[2 + 4D_\Theta/D + 2(D_\Theta/D)^2 - D_\Theta/D - 2(D_\Theta/D)^2 - (D_\Theta/D)^3 - 2 + 4D_\Theta/D - 2(D_\Theta/D)^2$$

$$- D_\Theta/D + 2(D_\Theta/D)^2 - (D_\Theta/D)^3] - \Theta D_\Theta$$

$$= (cD^2/4)[6D_\Theta/D - 2(D_\Theta/D)^3] - \Theta D_\Theta$$

$$= (cDD_\Theta/2)(3 - (D_\Theta/D)^2) - \Theta D_\Theta.$$

Again using (4), we have

$$S_\Theta = (cDD_\Theta/2)(3 - 1 + 2\Theta/3Dc) - \Theta D_\Theta$$

$$= cD_\Theta(D + \Theta/3c) - \Theta D_\Theta. \tag{6}$$

From (4), isolating D, we have $D^2 - 2\Theta D/3c - {D_\Theta}^2 = 0$. Solving for $D$, we find

$$D = \Theta/3c + \sqrt{{D_\Theta}^2 + (\Theta/3c)^2}, \tag{7}$$

from which we exclude the negative root given our constraints ($D \geq 0$, $D_\Theta \geq 0$, $0 \leq \Theta \leq 3Dc/2$).

Applying (7) into (6), we finally arrive at

$$S_\Theta = cD_\Theta\left(\sqrt{{D_\Theta}^2 + (\Theta/3c)^2} + 2\Theta/3c\right) - \Theta D_\Theta. \tag{8}$$

In the definition of a hard threshold, $S_\Theta^H$ adds the $\Theta \times D_\Theta$ rectangle below the threshold (Suppl. Fig. 4) and obtains the size scaling

$$S_\Theta^H = cD_\Theta\left(\sqrt{{D_\Theta}^2 + (\Theta/3c)^2} + 2\Theta/3c\right). \tag{9}$$

Given that any thresholding will affect scaling estimates, we note that (8) provides a higher estimate of the scaling exponent $\chi_{sh}$, whereas (9) provides the lower estimate. Generally, the error introduced by thresholding will be small if the threshold is small with respect to the amplitude of the parabola. Accordingly, in our analysis, we set our initial threshold Θ for $k$ = 1 small, i.e., at -2SD.

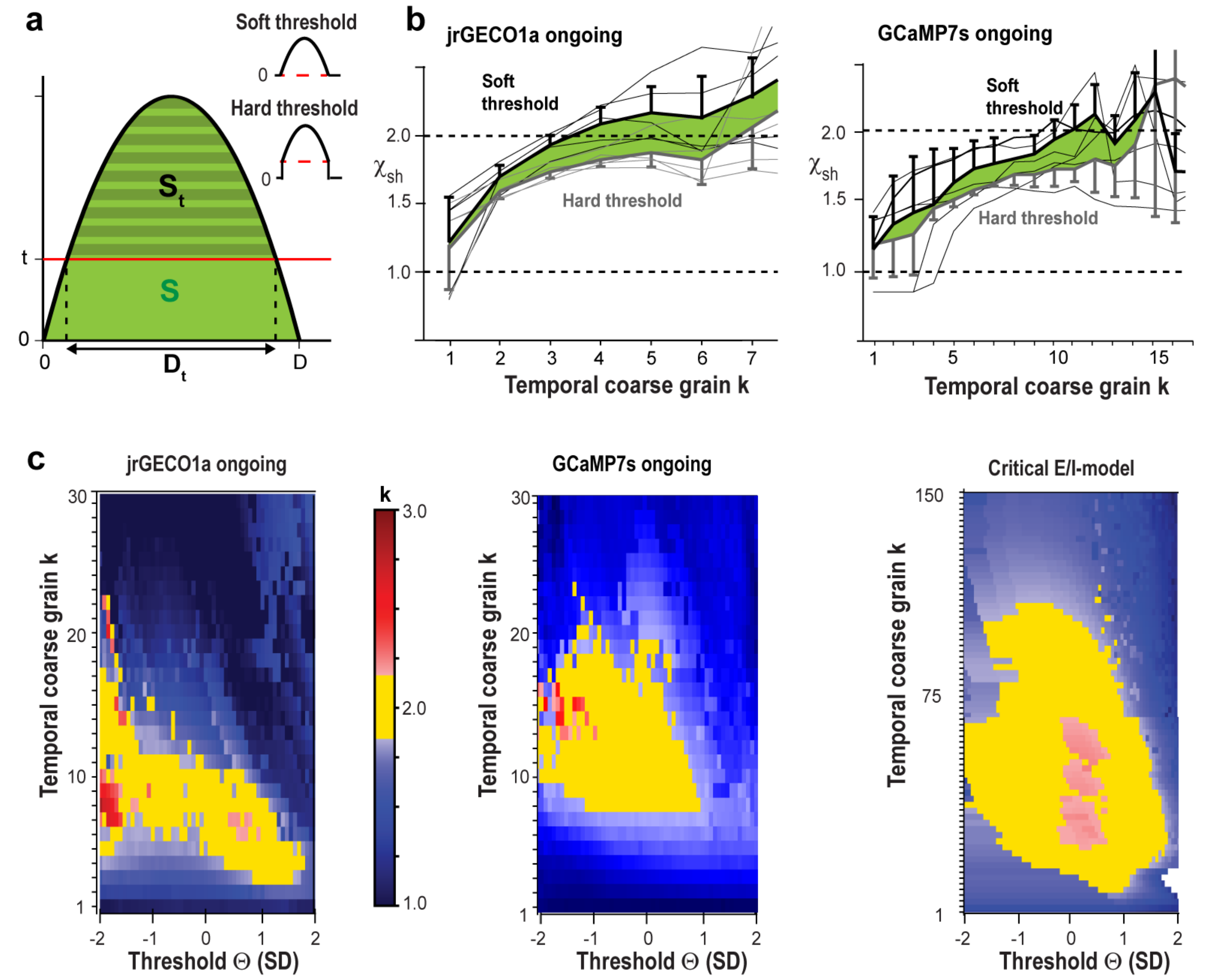


**Supplementary Figure S6 | Robust estimates of the scaling exponent $\chi_{sh}$ = 2 for two types of thresholding and wide ranges of threshold Θ**. (**a**) Sketch depicting the derivation of the scaling error as a function of threshold Θ. *Insets*: Soft thresholding subtracts the threshold from the time course ($S_t$), whereas in hard thresholding, all subthreshold values are set to 0 and *S* is maintained. (**b**). Soft (*black*) and hard (*grey*) thresholding provides an upper and lower error estimate of $\chi_{sh}$ respectively (*green area*). *Left*: jrGECO1a (n = 5 mice; ongoing activity). *Right*: GCaMP7s (n = 3 mice; ongoing activity). *Thin lines*: single animals. *Thick lines*: average. (**c**) Color maps depicting the value of $\chi_{sh}$ as a function of *Θ* and *k*. Note that $\chi_{sh}$ remains close to 2 for a wide range of Θ(-2) to Θ(0) covering an increase of avalanches by an order of magnitude (*yellow region*). *Left*: Average over n = 5 mice (jrGECO1a). *Middle*: Robust scaling estimates are already found at the level of a single mouse (GCaMP7s). *Right*: Results for our critical E/I-model (g = 3.5).

#### Supplementary Figure S7

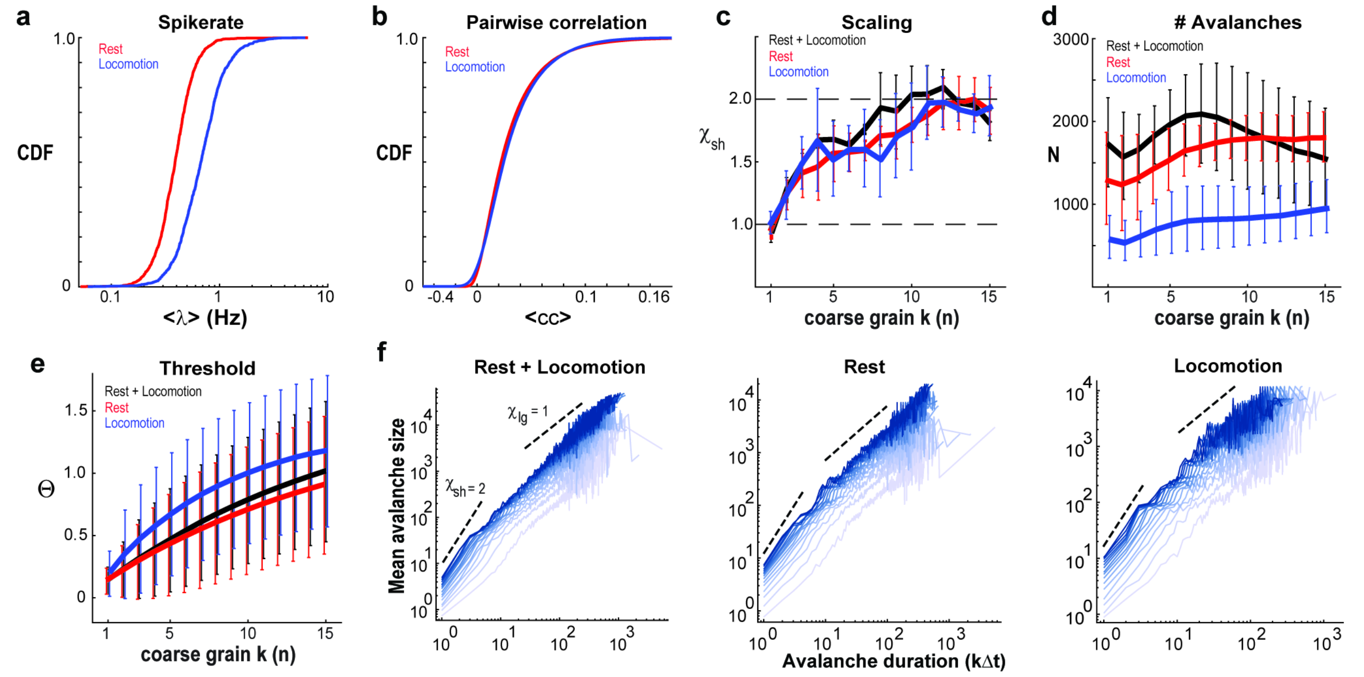


**Supplementary Figure S7 | Robustness of parabolic avalanches in mPFC/ACC to changes in behavioral states such as quiet rest vs. locomotion**. (**a**) Spike rate and pairwise correlation cumulative density functions for periods of locomotion and quiet rest. Spike rate distributions differ significantly between behavior states. (**c**) Scaling exponent vs. coarse grain factor for resting, locomotion, and resting + locomotion. (**d**, **e**) Number of avalanches and threshold value as a function of coarse grain factor *k*. (**f**) Mean avalanche size vs. duration curves for individual coarse grain factors. Due to poor statistics, scaling curves for avalanches during locomotion are noisy in relation to avalanches seen at rest. This noise is more apparent when attempting to obtain fits for scaling exponents (panel c).

#### Supplementary Figure S8

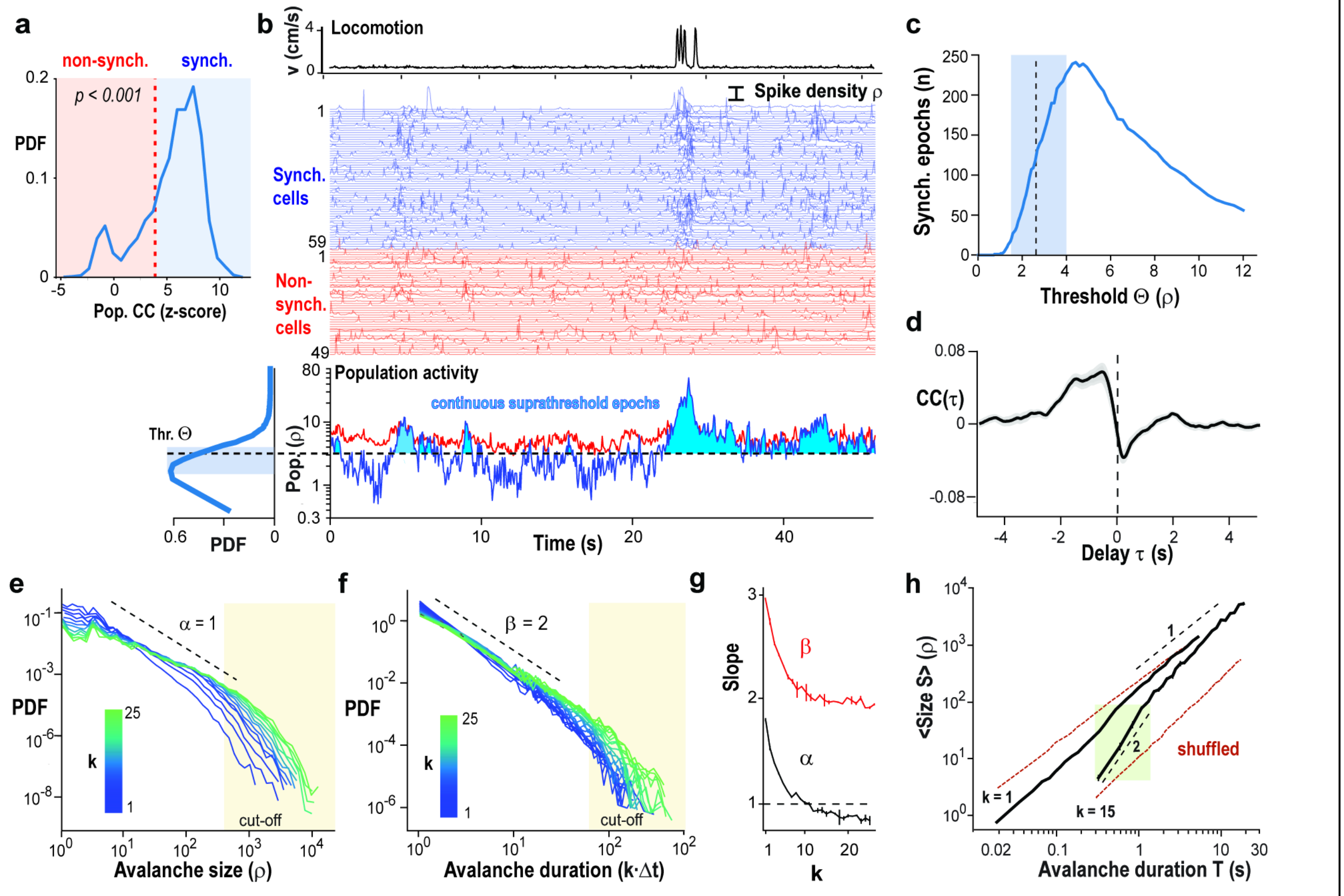


**Supplementary Figure S8 | Neuronal synchronization during ongoing activity in prefrontal cortex measured using 2PI with GCaMP7s.** (**a**) A minimal z-score (*p < 0.001; broken line*) in the correlation with population activity was used to separate synchronized (*blue*) from non-synchronized neurons (*red*). (30 min recording, single mouse). (**b**) Wheel locomotion (*top*), spike density raster separated into synchronized (*blue*) and non-synchronized neurons (*red*) and corresponding population activity, the summed spike densities per time bin *Δt* (*bottom*). Population activity above threshold Θ (*broken line*) establishes epochs of continuous suprathreshold activity (*filled blue areas*). Note reduced fluctuation in population activity of non-synchronized neurons. *Bottom left*: Distribution of population activity. Ongoing activity in mPFC/ACC of a single mouse using prism-based 2PI (See also Suppl. Fig. S1). (**c**) Number of epochs as a function of absolute threshold Θ (Single mouse; linear scale). (**d**) Locomotion onset is preceded by mPFC/ACC activity. Cross correlation (*CC*) between locomotion velocity and population activity (average from n = 3 mice). (**e**) Power law distributions in the size of suprathreshold epochs identifies the hallmark of neuronal avalanches. Distributions become shallower with temporal coarse graining *k* (*Δt* = 22 ms; 45.5 . Note cut-off region for sizes *S* >~500. (**f**) Corresponding power law distributions in epoch duration and cut-off region starting approximately for *L* >~80. (**g**), Change in the slope for size (α) and duration (β) distribution with temporal coarse graining *k*. Note the crossing of α = 1 at *k* ≅ 10. (**h**) $\chi_{sh} \cong 2$ holds for avalanches of duration 0.3 – 1.7 s at *k* = 15 (*green region*), not found at *k* = 1 and corresponding shuffled conditions (*red*).

#### Supplementary Figure S9

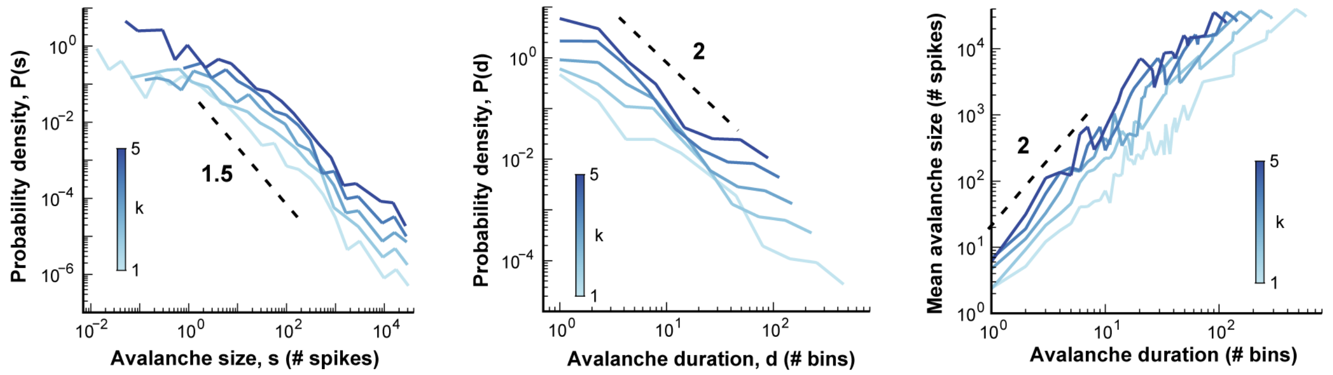


**Supplementary Figure S9 | Example of avalanche size, duration distributions and scaling from large field 2-PI using a mesoscope.** (**a**) Distributions for avalanche sizes (*left*), durations (*middle*), and scaling curves (*right*) obtained using different temporal coarse graining $k$ ($\Delta t$ = 100 ms; 10 ; mouse 1). Distributions become shallower with increase in $k$.

#### Supplementary Figure S10

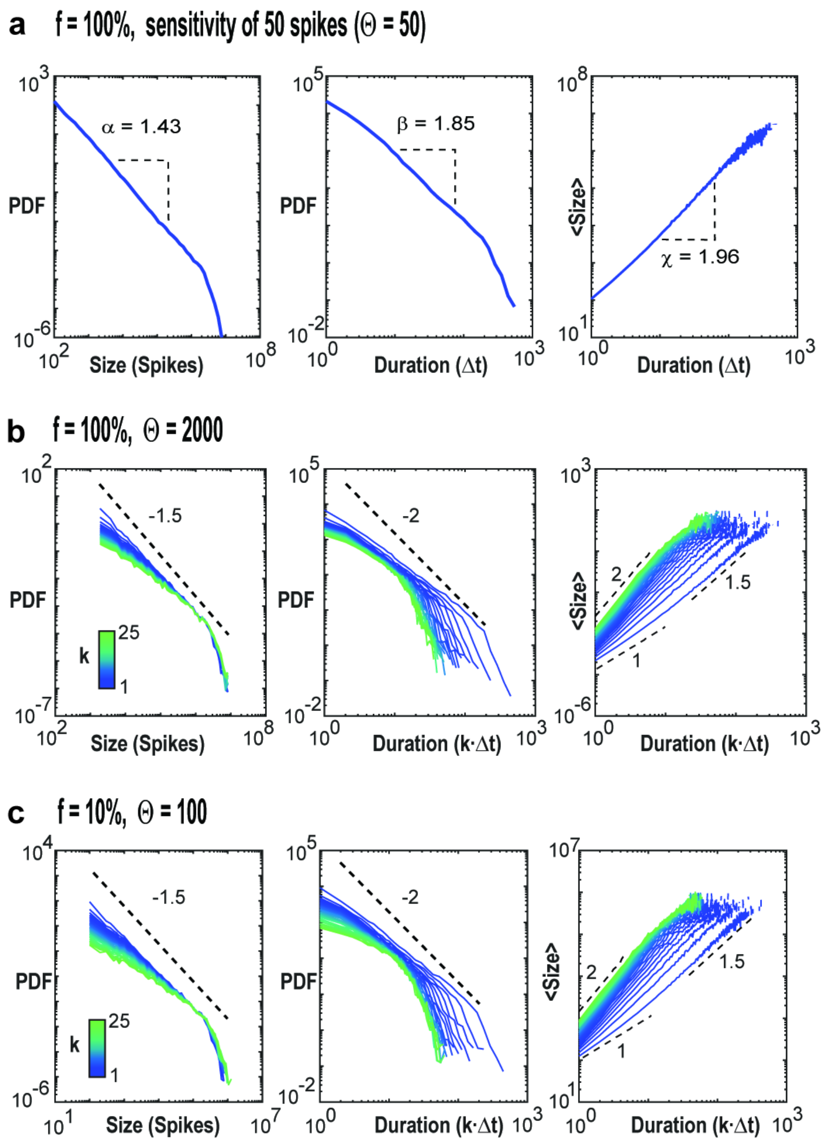


**Supplementary Figure S10| Examples of avalanche distributions as a function of threshold *Θ* and sampling fraction *f* for critical E/I-model.** (**a**) Distributions for the fully sampled model (*P* = 1; N = $10^6$ neurons, g = 3.5; $10^8$ time steps) at population spike sensitivity of 50 spikes/time step (*Θ* = 50). *Left*: Size. *Middle*: Duration. *Right*: Mean size vs. duration scaling with $\chi = 1.96$. Note that the Kinouchi model requires about a minimum of 100 spikes in N = $10^6$ to initiate successful propagation. The fully sampled model exhibits critical exponents close to $\alpha = 3/2$, $\beta = 2$ that fulfill the analytical prediction of $\chi = (\beta\text{-}1)/(\alpha\text{-}1) = 2$. (**b**) Temporal coarse graining recovers $\chi \approx 2$ at low sensitivity, i.e., high *Θ*. Example distributions and scaling for the fully sampled model (*f* = 100%) and high threshold *Θ* = 630. (**c**) As in b with 10 times reduced sampling *f* = 10% and 20 times reduced threshold *Θ* = 100. For corresponding χ(k) plots see main text Fig. 3. Note that in this model, uncorrelated propagated activity yields long-duration avalanches with $\chi_{lg} = 1.5$. T = $10^8$ time steps. *Broken, black lines*: slopes as visual guide to the eye.

78 **Supplementary Figure S11**

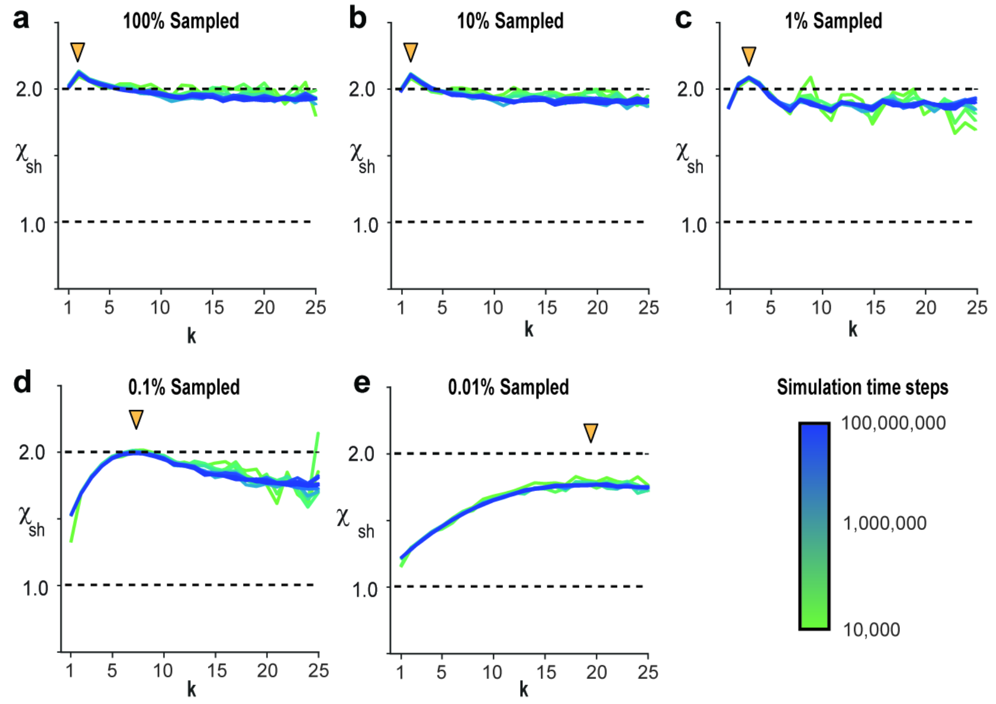


**Supplementary Figure S11 | Short simulation times provide robust estimates of $\chi_{sh}$ as a function of temporal coarse-graining *k* under fully-sampled and subsampled conditions.** (**a**) In the fully sampled model (*f* = 100%), reasonable estimates of $\chi_{sh}$(k) are obtained with 10,000 simulation time steps that only marginally improve even for 10,000 times longer simulations. (***b*** – ***e***) Same as in *a* but for increasingly lower sampling fractions *f*. For increasingly lower sampling (*f* = 10 – 0.1%), recovery of $\chi_{sh}$ = 2 requires a corresponding increase in temporal coarse-graining factor *k* until recovery fails for very low sampling fractions (*f* = 0.01%; *arrow heads*).

79 **Supplementary Figure S12**

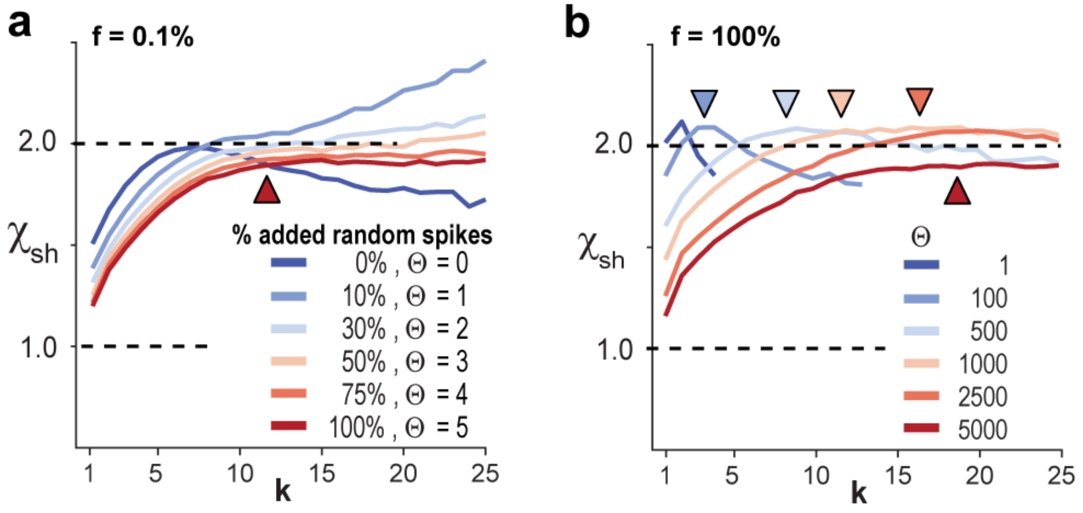


**Supplementary Figure S12| Impact of added irregular firing and subsampling as a function of threshold on the recovery of $\chi_{sh}$.** (**a**) Increasing percentage of irregular spikes in the subsampled model ($f$ = 0.1%) necessitates a decrease in sensitivity, i.e., higher $\Theta$, to offset the effect of increased noise. This increase in $\Theta$ shifts the recovery of $\chi(k) \cong 2$ towards larger k and this recovery begins to fail ($\chi_{sh}(k) < 2$) at higher percentages of irregular firing (*arrowhead*). (**b**) Increased sensitivity, i.e., lower $\Theta$, in the fully sampled model ($f$ = 0.1%) shifts the recovery of $\chi(k) \cong 2$ towards smaller $k$ (*arrow heads*) as observed with deep-IP. Note failure of recovery when $\Theta$ is too high, for which $\chi_{sh}(k) < 2$ remains below 2 ($f$ = 100%). N = $10^6$ neurons, T = $10^8$ time steps.

#### Supplementary Figure S13

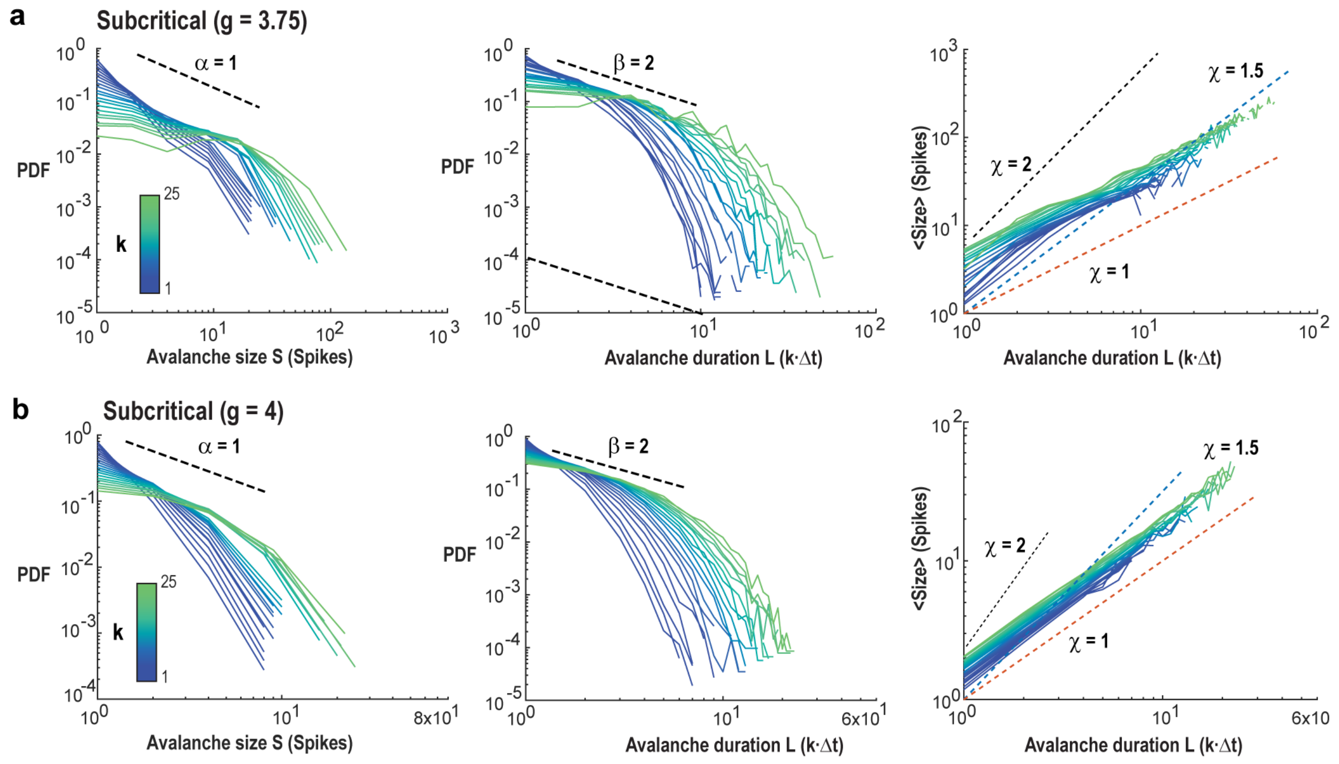


**Supplementary Figure S13 | Avalanche distributions and scaling as a function of temporal coarse graining *k* for subcritical network dynamics.** **(a)** (Distributions and scaling for subcritical dynamics (increased inhibition; g = 3.75). *Left*: Size. *Middle*: Duration. *Right*: Mean size vs. duration scaling with $\chi$ = 1, 1.5, 2. (**b**) Distributions and scaling for subcritical dynamics (increased inhibition; g = 4). Note failure of coarse graining to reach $\chi = 2$ for short or long duration avalanches. Instead, $\chi$ remains close to 1 for all avalanches. For corresponding $\chi$(k) plots see main text Fig. 2. For all simulations: $f$ = 0.1%; N = $10^6$ neurons, T = $10^8$ time steps. *Broken, black lines*: slopes as guide to the eye with corresponding slope values.

#### Supplementary Figure S14

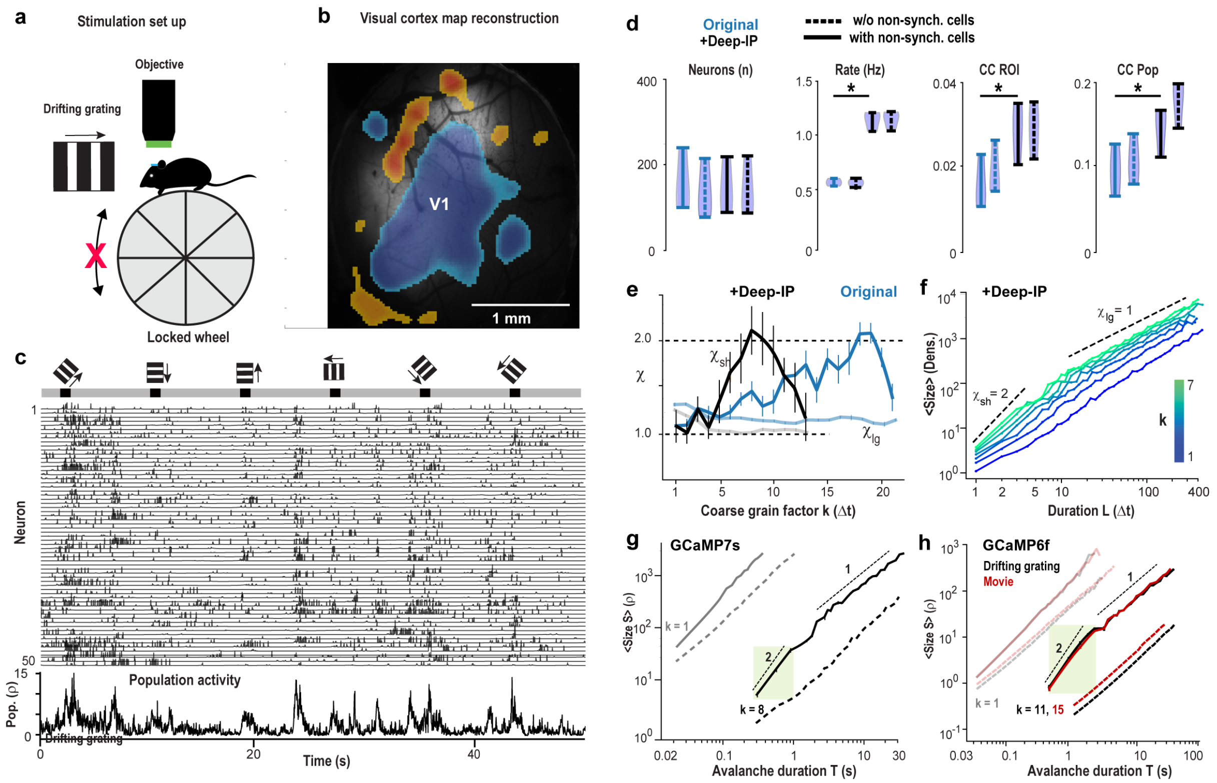


**Supplementary Figure S14 | Synchronized neurons exhibit avalanche scaling of $\chi_{sh}$ = 2 during visually evoked activity in V1**. (**a**) Sketch of 2PI during passive viewing of large-field gratings (8 directions) in head-fixed mice on a stationary wheel. (**b**) Example of a functional V1 map overlaid on the corresponding cortex surface (*single mouse*). (**c**) Example raster plot evoked by large field gratings (*top*). Note synchronized 'columnar' population responses to visual stimulation and during ongoing activity (+Deep-IP). *Bottom*: Corresponding population activity (Summed spike density, single mouse). (**d**) Deep-IP does not change the number of neurons identified (*Neurons*), but significantly increases spike rate (*Rate*), the pairwise cross-correlation among neurons (*CC ROI*) as well as the correlation between a single neuron and the remainder of the population (*CC Pop*). Note that evoked responses exhibited only a small, insignificant number of non-synchronized neurons (*broken lines*). (See also Suppl. Fig. S2 for comparison). (**e**) Summary of $\chi_{sh}$ and $\chi_{lg}$ as a function of $k$ for original (*Original*) and Deep-IP (+Deep-IP) processed neuronal activity. Note the shift to higher temporal resolution (smaller $k$) with +Deep-IP. (**f**) Mean-size vs. duration plot for $k = 1 - 7$ of evoked responses (+Deep-IP). Note $\chi_{sh}$ = 2 for short-lasting avalanches for $k$ ~7 ($k = 1 - 7$ shown) and the value of $\chi_{lg} \cong 1$ with temporal coarse graining. (**g**) $\chi_{sh}$ = 2 holds for evoked avalanches of duration T ~0.3 – 1.3 s at $k = 8$ (*green region*), which is abolished by trial-shuffling (*black broken line*). Corresponding $k = 1$ shown for comparison (*grey*). (**h**) Recovery of $\chi_{sh}$ = 2 in V1 during viewing of drifting gratings (*black*) or movies (*red*; 33 Hz, GCaMP6f; n = 6 mice; Allen Institute data set). $\chi_{sh}$ = 2 holds for avalanche durations ~0.3 – 3 s at $k$ = 11,15 (*green region*), which is abolished by trial-shuffling (*broken lines*). Corresponding $k = 1$ shown for comparison. *d* – *g*; Average from n = 3 mice.

**Supplementary Figure S15**

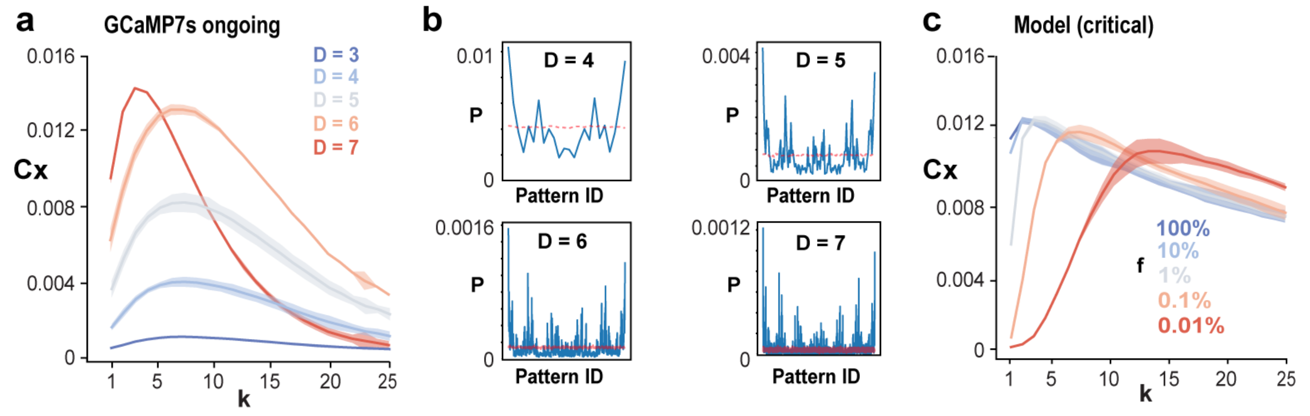


**Supplementary Figure S15| Peak in pattern complexity with temporal coarse graining is robust to pattern depth and threshold.** (**a**) Pattern complexity as a function of temporal coarse graining *k* for different pattern depths *D*. Ongoing activity in ACC/PFC measured with GCaMP7s (n = 3 mice; synchronized cells only, *cf*. Fig. 1). (**b**) Corresponding probability distribution of identified patterns (*Pattern ID*) for *D* = 4 – 7. (**c**) Subsampling shifts peak pattern complexity to higher coarse graining *k* (critical E/I-model).

85 **Supplementary Figure S16**

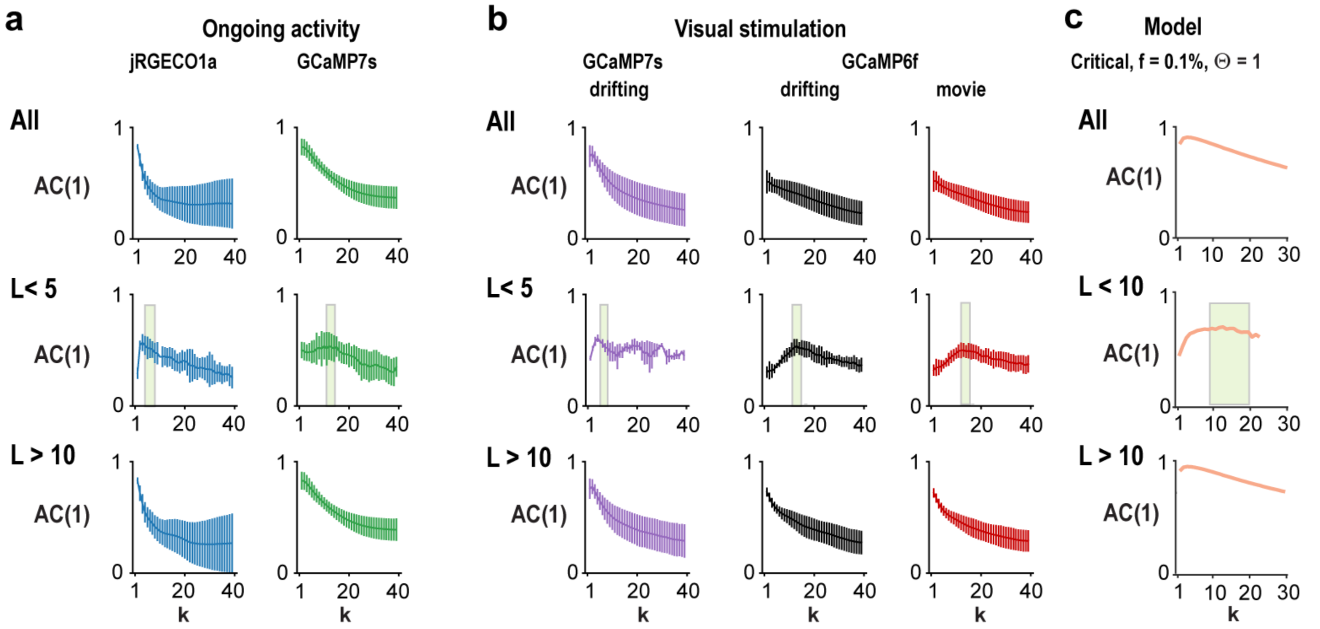


**Supplementary Figure S16 | Autocorrelation for short-lasting avalanches peaks at temporal coarse graining *k* that achieves $\chi_{sh}$ = 2.** (**a**) Change in autocorrelation (*AC(1)*) with temporal coarse graining *k* separated into all avalanches (*All*), short-lasting avalanches (*L* < *5*) and long-lasting avalanches (*L* > *10*). For short-lasting avalanches, the AC(1) peaks close to temporal coarse graining values at which the corresponding $\chi_{sh} \geq 2$ (*green area*). This relationship is not present for many-generation avalanches or when all avalanches are considered. (**b**) Corresponding analysis for visually evoked avalanches. (**c**) Corresponding analysis for the model with critical dynamics. Color code refers to Fig. 3d.

86

#### Supplementary Figure S17

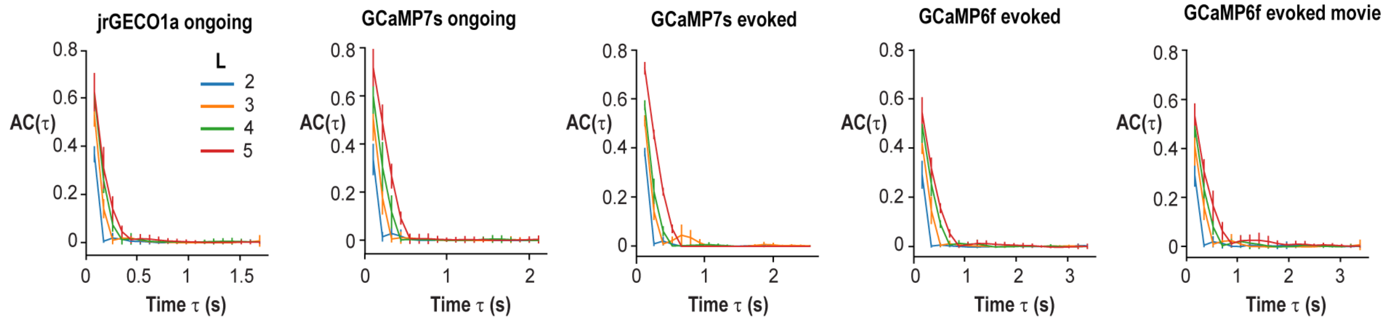


**Supplementary Figure S17 | Parabolic avalanches do not show periodic recurrence.** Autocorrelation for parabolic avalanches at temporal resolution for which $\chi_{sh} = 2$ separated by number of generations *L* (average over all mice per experimental condition). Note absence of recurrence supporting a non-oscillatory origin of parabolic avalanches. Mean ± SD.

**Supplementary Figure S18**

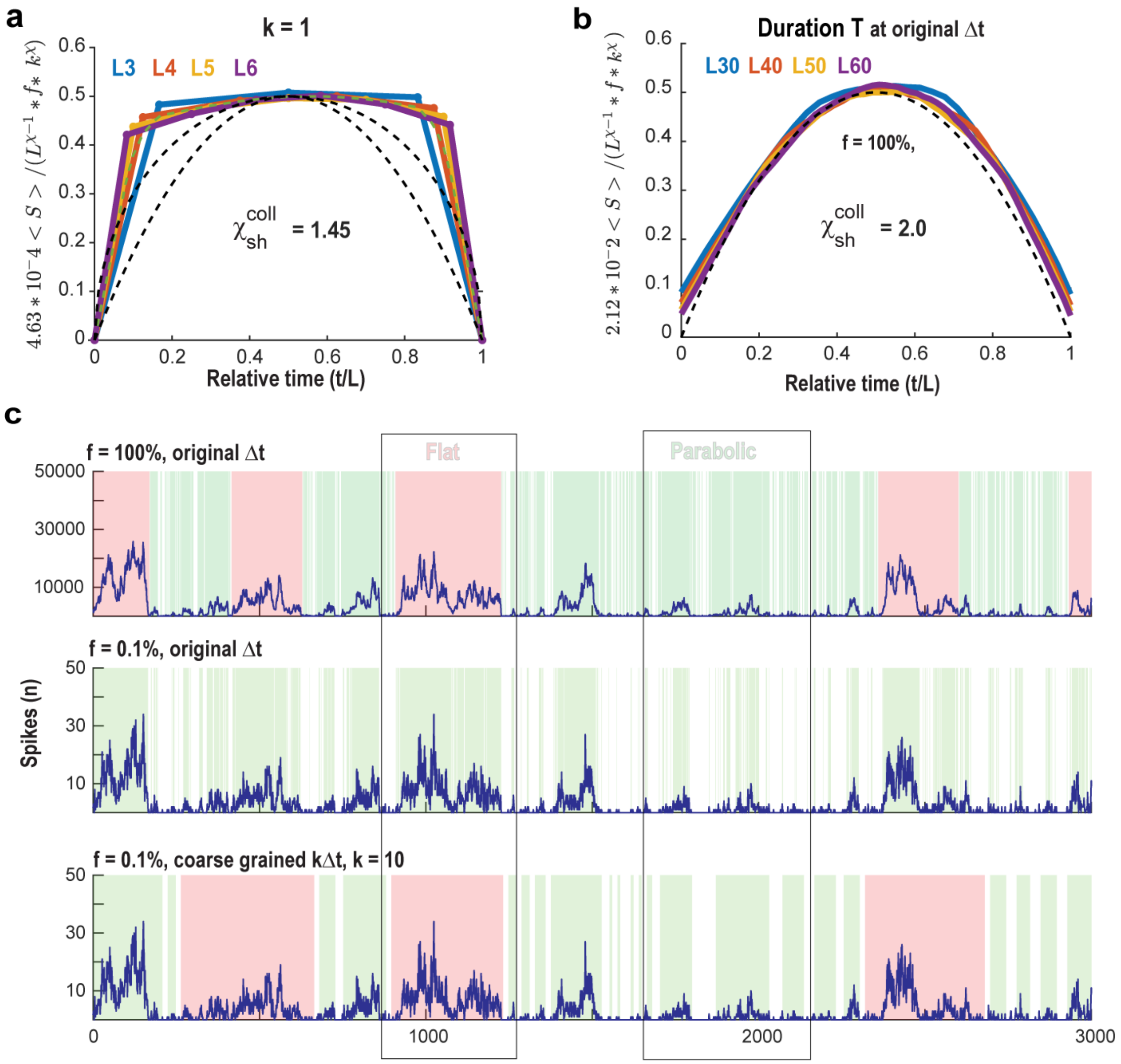


**Supplementary Figure S18 | Coarse-grained, few-generation avalanches with $\chi_{sh}$ = 2 exhibit inverted parabolic shape in cortical synchronization at high-resolution over many decades of generation**. (**a**) Critical E/I-model at $k = 1$ and corresponding avalanche collapse for $L = 3 – 6$; see also Fig. 5a. (**b**) Critical E/I-model. As in Fig. 5e for the fully sampled model, i.e., $f = 100\%$. Same as in *a* for $L = 30, 40, 50, 60$ generation avalanches at high temporal resolution. Note $\chi_{sh}^{coll} = 2.0$ for all generations. (**c**) Recovery of the time course in flat and parabolic avalanches at f=0.1% and temporal coarse graining of k = 10.
